# Histone H2B isoform *H2bc27* is expressed in the developing brain of mouse embryos

**DOI:** 10.1101/2024.05.28.596135

**Authors:** Saki Egashira, Kazumitsu Maehara, Kaori Tanaka, Mako Nakamura, Tatsuya Takemoto, Yasuyuki Ohkawa, Akihito Harada

## Abstract

Histones bind directly to DNA and play a role in regulating gene expression mediated by chromatin structure. DNA sequences of these histone genes are quite similar, which has hindered individual analyses. The exact function of the 13 different isoforms of histone H2B remains unclear. In this report, a comprehensive gene expression analysis of the H2B isoforms, focusing on tissue specificity, was conducted. We generated mice lacking the H2bc27 gene, which exhibited brain-specific expression of E14.5, and proceeded to characterize the brain tissue. The phenotype of *H2bc27* knockout brains was similar to that of wild-type brains, yet transcriptome analysis indicated that *H2bc27* is associated with regulating the expression of several functional genes involved in mouse brain development. The methods used in this study may serve to facilitate comprehensive H2B isoform analysis.

## Introduction

Organogenesis in multicellular organisms involves integrating cells that have acquired specific functions through differentiation during development. Cell differentiation is governed by the expression of genes, which in turn is driven by altering the chromatin structure into the relaxed or condensed form. The chromatin structure is a series of nucleosomes consisting of core histones, i.e., H2A, H2B, H3 and H4, with two molecules each forming an octamer, around which 145–147 bp of DNA is wrapped [1]. Therefore, the properties of histones likely affect the structure of chromatin.

There are several post-translational modifications of histones, including acetylation, methylation and phosphorylation, which occur in amino acid residues located in the N- and C-terminal tail regions of histones [2]. These histone modifications are referred to as the “histone code” hypothesis, and combinations of histone post-translational modifications are known to affect gene expression [3]. In addition to histone modifications, histone variants with slightly different amino acid sequences have also been shown to affect chromatin structure, thereby influencing gene expression [4]. For example, histone variants H2A.Z and H3.3 are enriched in active regulatory elements and contribute to activating transcription [5,6]. Furthermore, histone variants have been reported to play pivotal roles in specific cell types, such as H3mm7, which functions in the transcriptional regulation of mouse skeletal muscle differentiation, and H3.5, which is specifically expressed in primate testis [7,8]. Thus, the diversity of core histones adds greater complexity to regulating gene expression by chromatin dynamics.

Most histone gene variants are expressed in a replication-independent manner, as observed for H2A.Z and H3.3 genes [9]. In contrast, canonical histone genes are expressed in a replication-dependent manner. The chromosomal localization of genes encoding canonical histones is usually within four distinct histone clusters, whereas histone variants are scattered outside these clusters [10,11]. Even among histone genes located within a cluster, there are some genes that yield slightly different amino acid sequences, which are referred to as isoforms from among variants [12]. In germ cells, Th2a, Th2b and H3t histone isoforms are expressed and functional [13–15]. Interestingly, it remains unknown if histone isoforms can specifically function in other tissues. H2B is the most common histone type with at least 13 isoforms recognized [16,17], and H2bc21/Hist2h2b/H2be therein has recently been reported to have profound effects in mouse brain olfactory systems [18,19]. Therefore, we aimed to elucidate tissue-specific functions of other H2B isoforms. Thus, in this report, tissue-specific expression of H2B genes was investigated by mRNA-seq analysis of comprehensive sets of tissues. We then focused on *H2bc27*, also known as *Hist3h2ba* or *H2bu2*, which exhibits characteristic expression patterns in the embryonic brain and revealed functional analysis of *H2bc27* using knockout (KO) mice.

## Results

### H2B isoform *H2bc27* is specifically expressed in the brain

We used RNA-seq data from various tissues to investigate the tissue specificity of H2B gene expression. Histone genes closely resemble each other in DNA sequence, with multiple copies of a gene or variants within each histone cluster. Identifying which read is derived from which gene remains impractical, preventing successful quantification of histone expression levels for each isoform or variant. Those reads corresponding to several genes are often excluded from the analysis as multi-reads. In this study, we used Gibbs sampling of the posterior distribution by salmon [20] to statistically estimate the expression levels of histone genes, which allowed us to consider the uncertainty of the mapping, including multi-reads. We estimated the expression levels of all mouse H2B genes in 25 tissues based on Cold Spring Harbor Lab (CSHL) long-read RNA-seq data for 100 bp paired-end reads (GSE36025) (figure 1*a*, figure S1).

**Figure 1.**
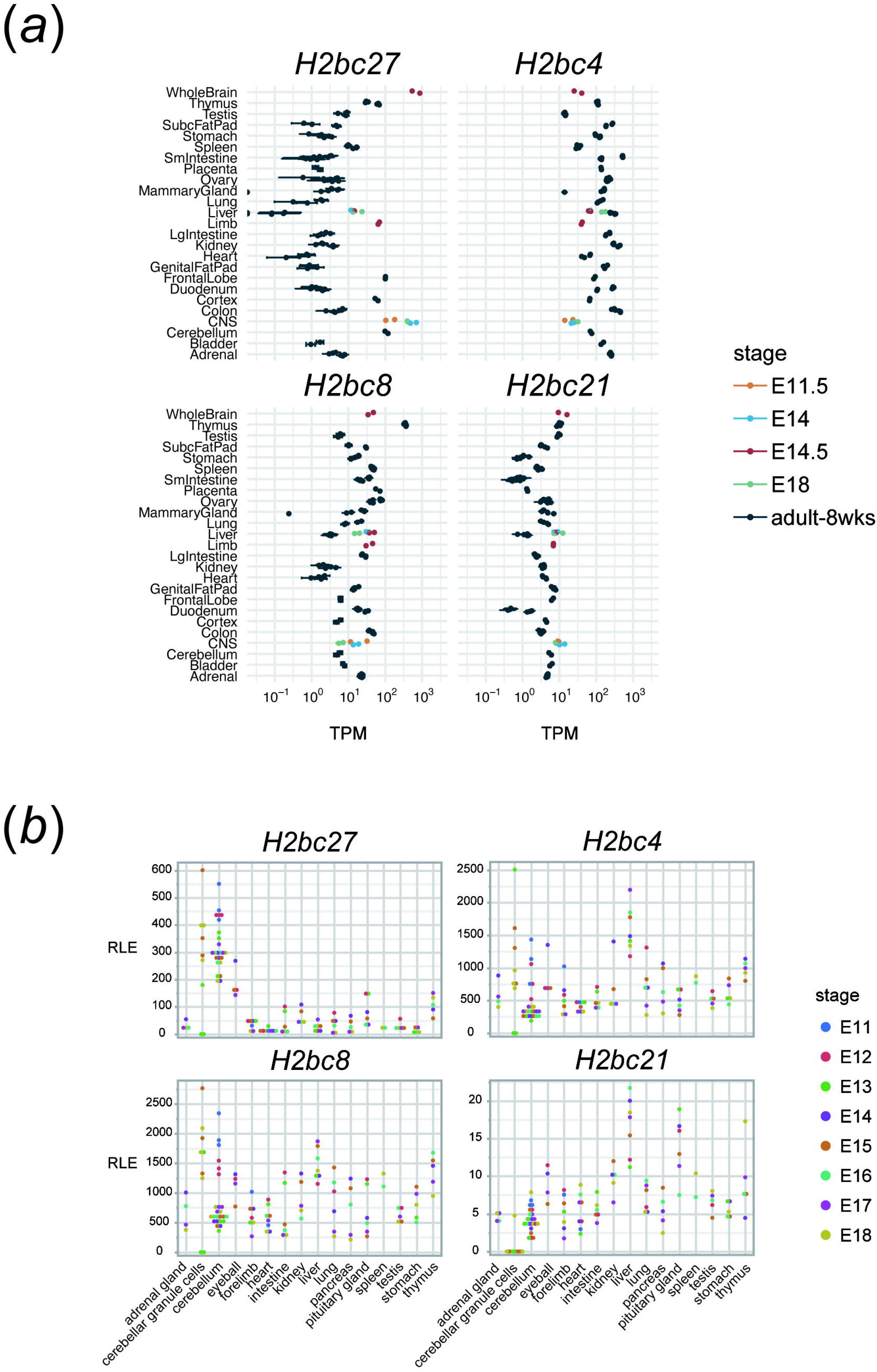
Analysis of H2B isoform gene expression for tissue specificity. *H2bc27* is expressed in a brain-specific manner. (*a*) Mouse H2B gene expression level analysis using Cold Spring Harbor Lab (CSHL) long-read RNA-seq data. The vertical axis indicates the tissue, and the horizontal axis indicates transcripts per million (TPM). (*b*) Mouse H2B genes expression level analysis using FANTOM5 CAGE-seq analysis. The vertical axis indicates relative long expression (RLE), and the horizontal axis indicates the tissues.

Comparison of the expression levels of each H2B histone between tissues revealed that *H2bc27* was highly expressed in whole brain and brain tissues (frontal lobe, cortex, CNS, and cerebellum) from embryonic stages (figure 1*a*, figure S1). Moreover, strong *H2bc1* and *H2bc22* expression was detected in testis, whereas high expression of *H2bc3, H2bc6, H2bc12, H2bc18*, and *H2bc23* as well as several others was observed in the thymus. However, evaluating the functional coherence of these H2B genes was not possible because they were found to be abundant in multiple tissues. Thus, we focused on *H2bc27* for the following analysis, whose expression was consistently high in the brain.

We analyzed CAGE-seq data published in the FANTOM5 (Functional ANnoTation Of the Mammalian genome) project to validate that *H2bc27* is expressed specifically in the brain. Of these, embryonic tissue data from the mouse promoterome dataset in ZENBU (http://fantom.gsc.riken.jp/zenbu) was selected, and the expression levels of *H2bc27* and several other H2B histones were confirmed. The results demonstrated that *H2bc27* was specifically expressed in brain tissues (cerebellum and cerebellar granule cells) during embryonic stages from E11 to E18 (figure 1*b*). Conversely, for other histones, expression was higher in the liver and pituitary gland, where the differences between these tissues and other tissues were minor or less specific.

Taken together, these findings indicated that *H2bc27* is specifically expressed at high levels in brain tissues from embryonic stages.

### *H2bc27* KO mouse brain presents a normal phenotype

We hypothesized that *H2bc27* is involved in brain development because this histone is specifically expressed in the brain during embryonic stages. We generated *H2bc27* KO mice to test this hypothesis using the CRISPR/Cas9 system [21]. As mentioned above, histone genes have similar DNA sequences. Thus, gRNAs were designed in the upstream and downstream regions with unique sequences outside the H2bc27 gene (figure 2*a*). The results of genotyping displayed a normal H2bc27 gene product of 2069 bp and a defective H2bc27 gene product of 985 bp, indicating that wild-type mice (*H2bc27*^+/+)^, KO mice (*H2bc27*^−/−^) and hetero KO mice with both (*H2bc27*^+/–^) were generated (figure 2*b*). Next, to evaluate brain morphology, we extracted the olfactory bulb to the pontine flexure of the brain at E14.5. No difference in the anatomical phenotype was identified among *H2bc27*^−/−^, *H2bc27*^+/–^, and *H2bc27*^+/+^ littermates (figure 2*c*). This observation suggests that *H2bc27* may not be important in brain organization during embryogenesis.

**Figure 2.**
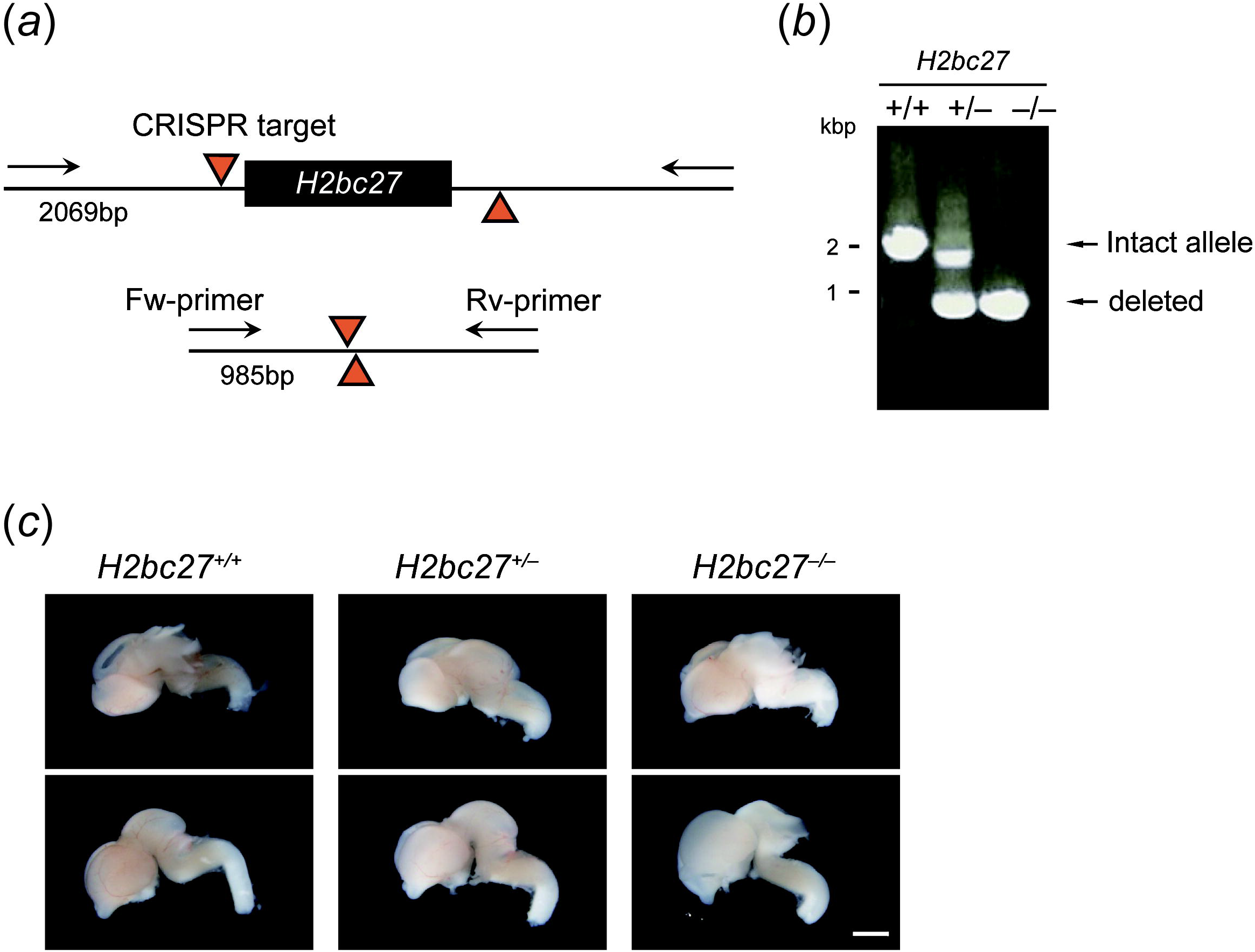
The *H2bc27* knockout (KO) system. (*a*) Strategy for constructing the *H2bc27* knockout using the CRISPR/Cas9 system. The gRNAs were designed with unique sequences in the upstream and downstream regions, i.e., outside the H2bc27 gene. (*b*) Agarose gel electrophoresis pattern for mouse genotyping. Wild-type, hetero KO and homo KO mice (referred to as *H2bc27*^+/–^, *H2bc27*^+/+^, and *H2bc27*^−/−^ respectively) were produced. (*c*) Representative images of two replicates of *H2bc27*^+/–^, *H2bc27*^+/+^, and *H2bc27*^−/−^ mice. There were no notable anatomical phenotype differences in the mice brains at E14. Scale bar, 2 mm.

### H*2bc27* is involved in regulating functional gene expression

The anatomical analysis of *H2bc27* KO mice did not provide direct evidence that *H2bc27* is required for brain morphogenesis in mouse embryos. Thus, RNA-seq analysis was carried out to determine whether gene expression in brain tissues was altered.

Total RNA-seq was performed on purified RNA from brains of E14 embryos derived from two heterozygous KO pregnant mice crossed with a male heterozygous mouse. Differentially Expressed Genes (DEGs) could not be extracted among *H2bc27*^−/−^, *H2bc27*^+/–^, and *H2bc27*^+/+^ embryos. Nonetheless, a comparison between *H2bc27*^−/−^ and *H2bc27*^+/+^ revealed that H2bc27 and Col4a1 gene expression were clearly downregulated in *H2bc27*^−/−^ embryos (Table 1). In relation to the brain, Col4a1 is associated with cerebral microvascular disease [22]; however, its relevance to brain development remains elusive.

**Table 1.** Result of a two-group comparison analysis of differential expressed genes between *H2bc27*^−/−^ and *H2bc27*^+/+^.

|  |  |  |
| --- | --- | --- |
| **ens_gene** | ENSMUSG00000056895 | ENSMUSG00000031502 |
| **ext_gene** | *H2bc27* | *Col4a1* |
| **biotype** | protein_coding | protein_coding |
| **chr** | 11 | 8 |
| **baseMean** | 125.0333142 | 1579.029747 |
| **homoKO_vs_wt** | -6.672165933 | -0.573633497 |

We further evaluated components that separate *H2bc27*^−/−^ from *H2bc27*^+/–^ and *H2bc27*^+/+^ by principal component analysis (PCA) and were able to distinguish these three phenotypes by the component of PC4 (6.7%) (figure 3*a*). Of the 500 genes attributed to PC4, 212 genes contribute to PC4(–), which is biased toward *H2bc27*^−/−^, and 288 genes contributed to PC4(+), which is biased toward *H2bc27*^+/+^. Next, to infer possible changes in brains of *H2bc27*^−/−^, gene ontology (GO) analysis was performed for genes contributing to each of the two groups. As a result, PC4(–) characterizing *H2bc27*^−/−^, was enriched with terms related to the following developmental events: brain development [GO: 0007420], neuron projection development [GO: 0031175] and sensory organ development [GO: 0007423]. PC4(+) characterizing *H2bc27*^+/+^ was enriched with terms related to head development [GO: 0060322] and protein digestion and absorption [mmu04974] (figure 3*b*).

**Figure 3.**
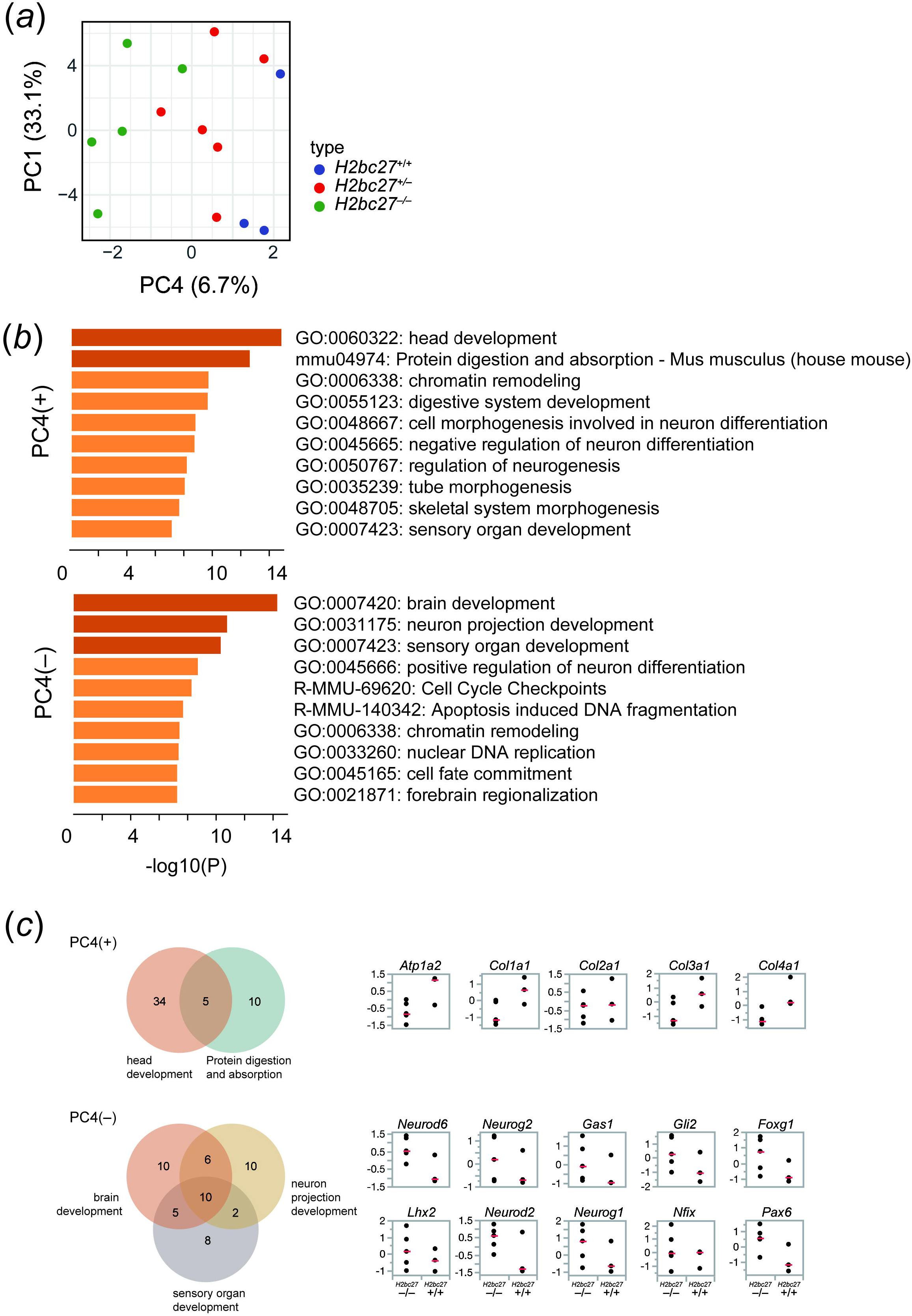
PCA and GO analyses to evaluate alterations associated with the *H2bc27* deletion. (*a*) PCA separated *H2bc27*^−/−^, *H2bc27*^+/–^, and *H2bc27*^+/+^ by the PC4 component. (*b*) GO analysis results for PC4(+) biased toward *H2bc27*^+/+^ vs. PC4(–) biased toward *H2bc27*^−/−^. The top 10 enriched terms for each are shown. (*c*) Expression levels by the *z*-score of genes overlapping with the top terms.

We then examined for bias of gene expression in these terms between *H2bc27*^+/+^ and *H2bc27*^−/−^ to further assess whether changes involving these terms were likely to be induced by association with *H2bc27*. Specifically, we extracted overlapping genes across the top terms as representatives, and their gene expression levels in each sample were compared by *z*-score analysis. The results indicated that ATPase, Na+/K+ transporting, alpha 2 polypeptide (*Atp1a2*), collagen, type I, alpha 1 (*Col1a1*), collagen, type II, alpha 1 (*Col2a1*), collagen, type III, alpha 1 (*Col3a1*), and collagen, type IV, alpha 1 (*Col4a1*), genes extracted from head development, protein digestion and absorption, as the top terms of PC4(+), tended to have lower expression levels in *H2bc27*^−/−^ embryos (figure 3*c*). In contrast, the PC4(–) overlapping genes for brain development, neuron projection development and sensory organ development, which are neurogenic differentiation 6 (*Neurod6*), neurogenin 2 (*Neurog2*), growth arrest specific 1 (*Gas1*), GLI-Kruppel family member GLOI2 (*Gli2*), forkhead box G1 (*Foxg1*), LIM homeobox protein 2 (*Lhx2*), neurogenic differentiation 2 (*Neurod2*), neurogenin 1 (*Neurog1*), nuclear factor I/X (*Nfix*), and paired box 6 (*Pax6*), were typically highly expressed in *H2bc27*^−/−^. These gene expression-level biases indicated that *H2bc27* may transcriptionally regulate these genes.

Collectively, these results suggest that *H2bc27* is associated with the expression of functional genes, i.e., *Neurod6, Neurog2, Gas1, Gli2, Foxg1, Lhx2, Neurod2, Neurog1, Nfix*, and *Pax6*, involved in the development of brain, neuron projection and sensory organs, as well as *Atp1a2, Col1a1, Col2a1, Col3a1*, and *Col4a1*, which are associated with head development and protein digestion and absorption, and thus may have roles in differentiation and functional acquisition to some extent.

## Discussion

As with canonical histones, genes encoding histone H2B isoforms, including *H2bc27*, are located within the histone clusters. Histone regulation of gene expression throughout the cell cycle is a feature shared by histone variants, and some of those histone variants perform tissue-specific functions [23]. Despite the relatively advanced functional analysis of the H2A and H3 histone variants, our understanding of how the H2B isoforms regulate gene expression and tissue formation remains underdeveloped. Concurrently, precise analysis of histone gene expression is challenging because their gene sequences are very similar.

In this study, we addressed this issue and analyzed the tissue-specific functions of H2B isoforms. We explored H2B isoforms expressed in a tissue-specific manner by mRNA-seq analysis of comprehensive sets of samples. We then focused on *H2bc27*/*Hist3h2ba*/*H2bu2*, a member of cluster 3, which exhibited a characteristic expression pattern in the embryonic brain, and attempted to analyze its function using KO mice.

Comprehensive expression analysis of H2B histone isoforms in various tissues revealed brain-specific expression of *H2bc27* from embryonic stages. The brain phenotype was not affected by the deletion of *H2bc27* at E14.5. For the gene expression analysis, GO analysis in PC4(–) indicated that brain, neuron projection and sensory organ development were enriched. Expression of genes included in these terms (e.g., *Neurod6, Neurog2, Gas1, Gli2, Foxg1, Lhx2, Neurod2, Neurog1, Nfix, Pax6*) increased in *H2bc27*^−/−^ mice. Consistent expression of these genes in a forebrain-specific manner suggests that *H2bc27* is involved in gene expression of forebrain differentiation (figure S2).

In summary, the results of this study indicated that although *H2bc27* has no significant effect on brain development, this H2B isoform may regulate the expression of several functional genes. Comprehensive characterization of all individual H2B isoforms to obtain tissue-specific functions should provide further insights into the role of H2B in development.

## Materials and Methods

### Animals

All animal procedures were conducted in accordance with the Guidelines for the Care and Use of Laboratory Animals and were approved by the Institutional Animal Care and Use Committee (IACUC) at Kyushu University.

### Generation of *H2bc27* KO mice and genotype analysis

*H2bc27* KO mice were generated with the CRISPR/Cas9 system, as described previously [21]. Briefly, mouse zygotes prepared by in vitro fertilization were electroporated with 0.1 μg/μl *Streptococcus pyogenes* Cas9 containing the nuclear localization sequence (NLS), 0.1 μg/μl crRNA and 0.1 μg/μl trans-activating CRISPR RNA (tracrRNA). The zygotes were cultured overnight and two-cell stage embryos were separated from unfertilized oocytes and damaged zygotes. The two-cell stage embryos were transferred into the oviduct of 0.5-dpc pseudopregnant recipient female mice. The following gRNAs targeting the upstream and downstream regions of the H2bc27 gene were used:

Upstream crRNA: 5’-CTGCCAATACACTTTCGGAT-3’

Downstream crRNA: 5’-TAATTAAGCAACAATCCCGC-3’

Genomic DNA was extracted from an ear punch. DNA was amplified by PCR with H2bc27 genotyping primers to detect wild-type and deletion alleles generated by genome editing. The PCR was performed with an initial incubation at 98 °C for 2 min, then 30–35 cycles of 98 °C for 10 s, 65 °C for 30 s and 68 °C for 120 s. The PCR product was detected by running a 1.0% (w/v) agarose gel with EtBr staining.

Primers used for genotyping were:

H2bc27 Forward: 5’-GTAATTATGGTACCCTTGCTTCAGC-3’

H2bc27 Reverse: 5’-GCTCAACTACATGTTCAAGACCAAT-3’

### Statistical inference of histone genes expression

We analyzed total of 125 samples of long RNA-seq from ENCODE/CSHL dataset (NCBI SRA: SRR453077-SRR453175, SRR567478-SRR567503), including 25 types of tissues and several developmental stages of mice (adult and E11.5-E18). To quantify the expression levels of histone genes along with the read assignment ambiguity (i.e., reads that fall within 2 or more genes), we used Salmon (version 0.12.0) [20] that enables sampling from posterior distribution of read counts on genes. The `salmon quant` was used with the options: --validateMappings -l A --numGibbsSamples 1000. The index was generated using GENCODE transcript reference (release M20).

### mRNA-seq and analysis

Following dissection at E14, the brain was removed from the olfactory bulb to the bridge flexure. After washing in PBS, brain tissues were transferred to 1.5 ml microcentrifuge tubes, and 500 μL of Trizol was added. Samples were homogenized with a pipette, incubated at 50 °C for 10 min, mixed thoroughly and stored at –30 °C. After RNA extraction, libraries were prepared using the SMART-Seq® Stranded Kit (TaKaRa Bio Inc. Shiga, Japan). The libraries were sequenced on NovaSeq6000 with cycles of 51+8+8+51.

Adaptor sequence and low quality sequence were removed and read length below 20bp were discarded by using Trim Galore. Reads were aligned to GRCm38 reference using HISAT2 and count matrix was obtained by using featureCounts.

Between-sample normalization was applied to the count matrix using median of ratios of geometric means [*] and principal component analysis was performed. Differentially expressed genes were extracted with DESeq2 using |log2FC| > 1 and padj < 0.1 as threshold values [24]. The gene ontology (GO) analysis was performed using Metascape (http://metascape.org) [25].

## Data accessibility

The datasets generated in this study are deposited in Gene Expression Omnibus under accession number (GSE268119).

## Acknowledgments

We are grateful to the staff of Ohkawa Lab. We thank Edanz (https://jp.edanz.com/ac) for editing a draft of this manuscript.

## Funding Information

This work was supported by JST CREST JPMJCR23N3 to K.M., JST PRESTO JPMJPR2026 to K.M., AMED BINDS JP22ama121017j0001 to Y.O., AMED-CREST JP23gm1810008 to A.H., JSPS KAKENHI JP21H05292, JP23K27087 and JP23H02394 to A.H., JP22H04696 and JP23H04288 to K.M., JP18H05528, JP24H02323, JP23H00372, JP22H04676 and JP22K19275 to Y.O. This work was also supported in part by the MEXT Promotion of Development of a Joint Usage / Research System Project :Pan-Omics DDRIC, MRCI for High Depth Omics, CURE:JPMXP1323015486 for MIB and RIIT in Kyushu Univ.

## Author disclosure statement

S.E.: Investigation, Validation, Visualization, Writing – original draft, Writing – review and editing; K.M.: Formal Analysis, Methodology, Software, Validation, Writing – original draft; K.T.: Data curation, Formal Analysis, Funding acquisition, Software; M.N.: Supervision, Writing – review and editing; T.T.: Investigation, Resources, Writing – review and editing; Y.O.: Conceptualization, Funding acquisition, Supervision, Writing – review and editing; A.H.: Conceptualization, Funding acquisition, Investigation, Project administration, Supervision, Writing – original draft, Writing – review and editing.

## Conflict of interest declaration

Authors declare no competing interests.

## Notes

### Competing Interest Statement

The authors have declared no competing interest.

