## Supplemental Materials for "Histone H2B isoform *H2bc27* is expressed in the developing brain of mouse embryos"

**Supplemental Material**

**Supplementary Figures 1-2 from Histone H2B isoform H2bc27 is expressed in the developing brain of mouse embryos**

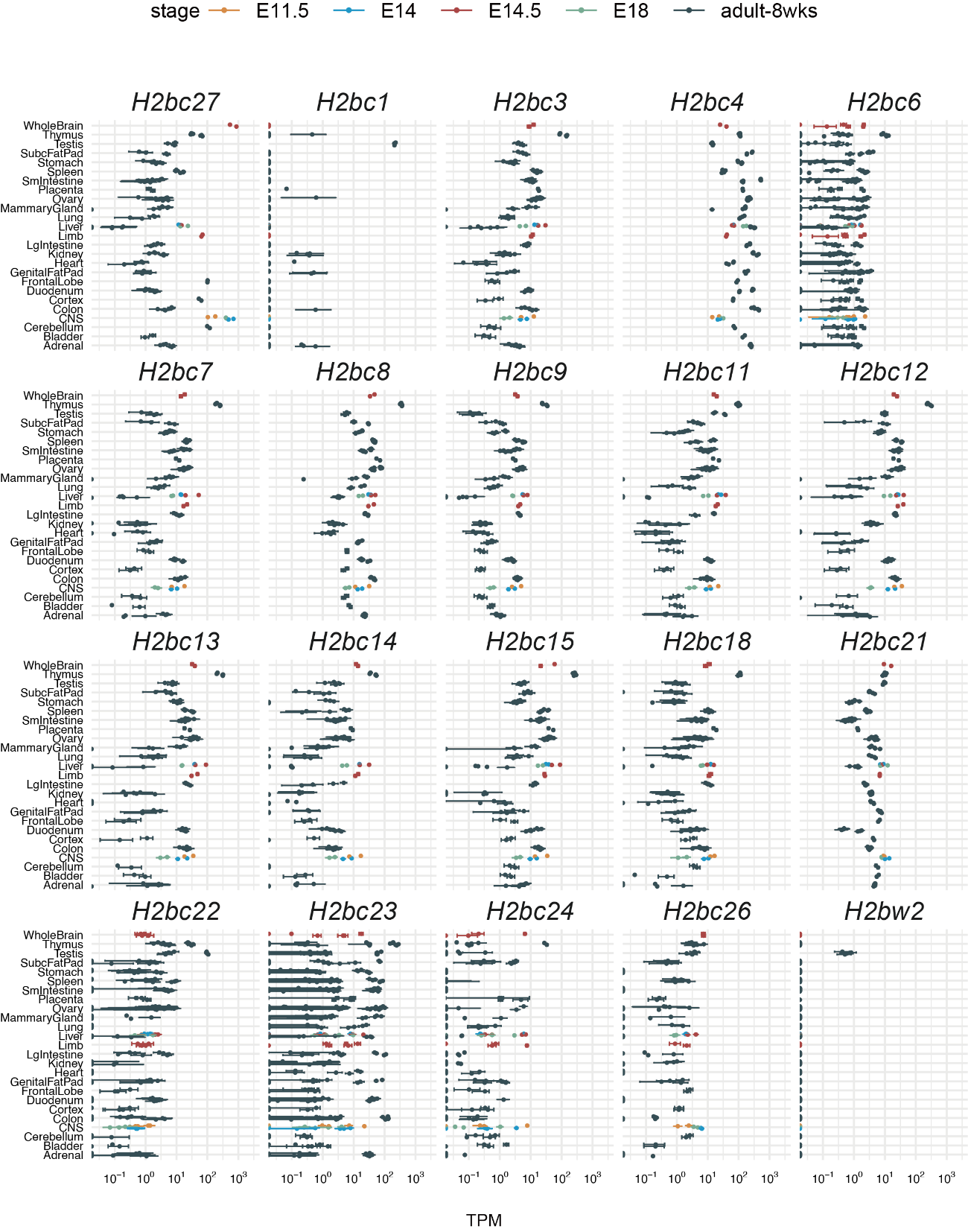

**Figure S1.** Analysis of all mouse H2B gene expression levels based on Cold Spring Harbor Lab (CSHL) long read RNA-seq data.

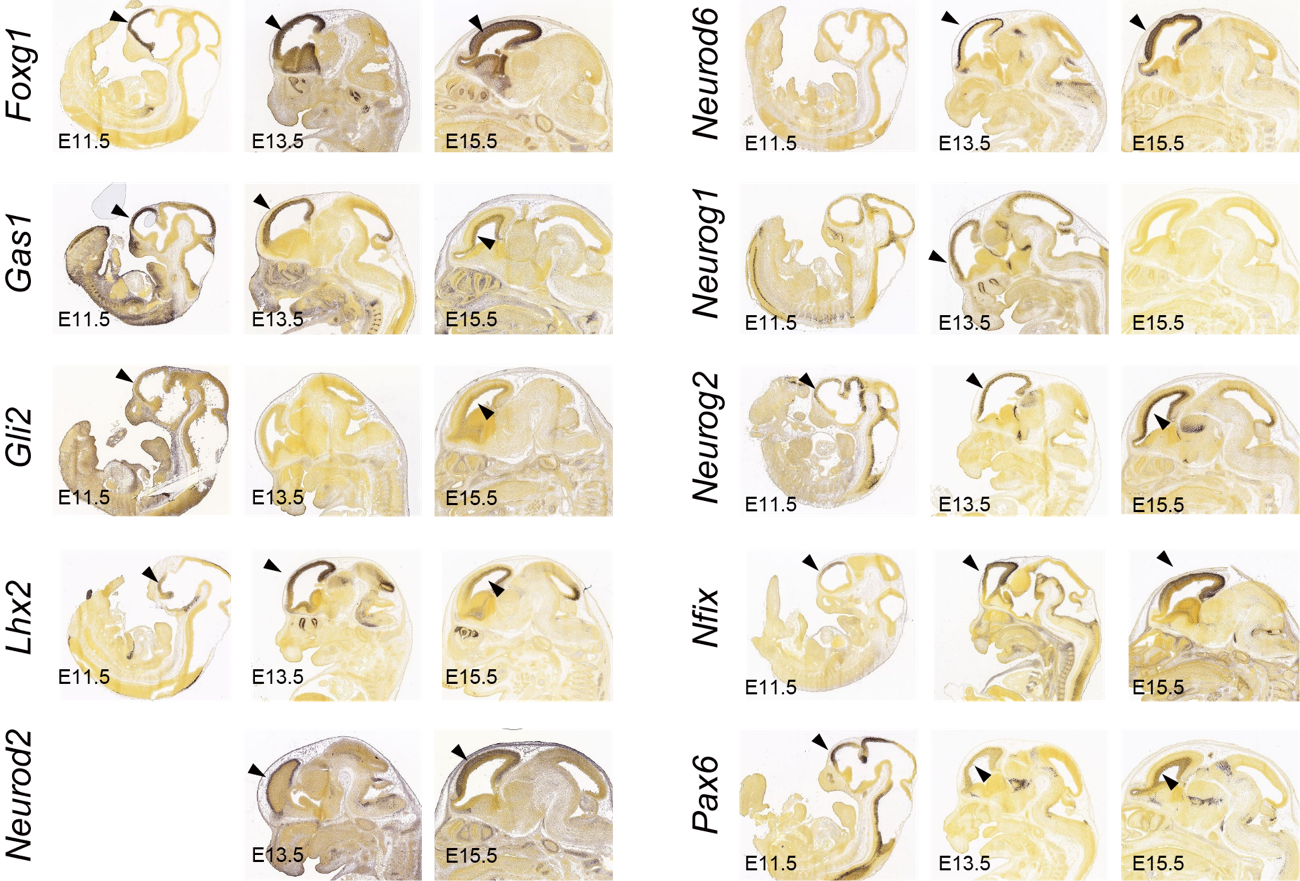

**Figure S2.** Developing mouse brain in situ hybridization (ISH) image data obtained from Allen Brain Atlas for overlapping genes in the top terms of PC4(+). Forebrain-specific localization is shown.
